# Synergism and antagonism of bacterial-viral co-infection in the upper respiratory tract

**DOI:** 10.1101/2020.11.11.378794

**Authors:** Sam Manna, Julie McAuley, Jonathan Jacobson, Cattram D. Nguyen, Md Ashik Ullah, Victoria Williamson, E. Kim Mulholland, Odilia Wijburg, Simon Phipps, Catherine Satzke

**Affiliations:** Infection and Immunity, Murdoch Children’s Research Institute, Royal Children’s Hospital, Parkville, Victoria, Australia; Department of Paediatrics, The University of Melbourne, Parkville, Victoria, Australia; Department of Microbiology and Immunology at the Peter Doherty Institute for Infection and Immunity, The University of Melbourne, Parkville, Victoria, Australia; Respiratory Immunology Laboratory, QIMR Berghofer Medical Research Institute, Herston, Queensland, Australia; Department of Infectious Disease Epidemiology, London School of Hygiene and Tropical Medicine, London, United Kingdom

**Keywords:** Streptococcus pneumoniae, pneumococcus, Respiratory Syncytial Virus, Pneumonia Virus of Mice, Murine Pneumonia Virus, Influenza, co-infection

## Abstract

*Streptococcus pneumoniae* (the pneumococcus) is a leading cause of pneumonia in children under five years old. Co-infection by pneumococci and respiratory viruses enhances disease severity. Little is known about pneumococcal co-infections with Respiratory Syncytial Virus (RSV). Here, we developed a novel infant mouse model of co-infection using Pneumonia Virus of Mice (PVM), a murine analogue of RSV, to examine the dynamics of co-infection in the upper respiratory tract, an anatomical niche that is essential for host-to-host transmission and progression to disease. Coinfection increased damage to the nasal tissue and increased production of the chemokine CCL3. Pneumococcal nasopharyngeal density and shedding in nasal secretions were increased by co-infection. In contrast, co-infection reduced PVM loads in the nasopharynx, an effect that was independent of pneumococcal strain and the order of infection. We showed this ‘antagonistic’ effect was abrogated using a pneumococcal mutant deficient in capsule production and incapable of nasopharyngeal carriage. The pneumococcal-mediated reduction in PVM loads was caused by accelerated viral clearance from the nasopharynx. Although these synergistic and antagonistic effects occurred with both wild-type pneumococcal strains used in this study, the magnitude of the effects was strain dependent. Lastly, we showed that pneumococci can also antagonize influenza virus. Taken together, our study has uncovered multiple novel facets of bacterial-viral co-infection. Our findings have important public health implications, including for bacterial and viral vaccination strategies in young children.

## INTRODUCTION

The severity and outcome of microbial infections are determined by a variety of host, pathogen, and environmental factors. As the pathogen colonizes the host, it encounters members of the resident microbiota and/or other pathogens. These interactions can influence microbial pathogenesis, including increased bacterial adhesion, enhanced virion stability, and modulation of the immune response by one microbe that benefits the other [1]. These interactions can be particularly relevant in anatomical sites that have complex microbial communities including the gastrointestinal and respiratory tracts [2–6].

Pneumococcus is a leading cause of community-acquired pneumonia in young children, particularly for those in low and middle-income settings [7,8]. This bacterium is a common resident of the upper respiratory tract in children. While nasopharyngeal colonization by pneumococci is usually asymptomatic, it serves as a prerequisite for disease and a reservoir for host-to-host transmission, which underpins herd protection conferred by pneumococcal conjugate vaccines [9,10]. Throughout history, 30-95% of severe or fatal cases during influenza pandemics were caused by secondary bacterial infections, with pneumococci being the leading etiologic agent [11]. In part, this is because the immune response to viral infection creates a hospitable environment for pneumococcal superinfection [11]. Influenza infection also has positive effects on pneumococci in the upper respiratory tract, exemplified by increased pneumococcal density in the nasopharynx of mice [12,13], as well as increased transmission to naïve hosts, which has been demonstrated in both animal and human studies [12,14,15].

Human Respiratory Syncytial Virus (RSV) disproportionately affects infants and is a major cause of bronchiolitis and infant hospitalization [2]. Although little is known regarding the mechanisms of pneumococcal-RSV co-infection, there is clear evidence of its clinical relevance. Detection of both RSV and pneumococcus in young children under two years of age presenting to hospital with acute respiratory infection is associated with increased disease severity [16]. Pneumococcal infection in pre-term infants hospitalized with RSV is associated with longer hospital stays and increased likelihood of admission to the intensive care unit [17]. Importantly, introduction of pneumococcal conjugate vaccination has been associated with a decline in RSV hospitalizations in children [18,19]. Lastly, direct interactions between the pneumococcal penicillin-binding protein 1a and RSV G protein enhances pneumococcal virulence in mice [20].

For this study, we developed an animal model to assess the dynamics of pneumococcal-RSV co-infection in the upper respiratory tract, which is currently poorly understood. Recent work has shown that most infants are already colonized by pneumococci at the time they contract RSV [21]. Additionally, children in low and middle-income settings are often colonized by pneumococci early in life [8] and are therefore also likely to carry pneumococci at the time of viral infection. To represent co-infections in these high-risk settings, we established a murine model where pneumococci colonize the nasopharynx before viral challenge. Human RSV replicates poorly in mice. Existing models therefore rely on administering high viral titers (~10^4^-10^6^ PFU) [22–26], which are less likely to be biologically relevant. This is because the infectious dose to establish RSV infection in adult volunteers is as low as 10^2^-10^3^ and is anticipated to be even lower in infants [27–29]. To overcome this, we used Pneumonia Virus of Mice (PVM, also known as Murine Pneumonia Virus), a murine analogue of RSV [30,31], and member of the *Orthopneumovirus* genus. PVM replicates in the airway epithelium and induces similar pathologic features in mice to that observed in human RSV disease [32,33]. By challenging pneumococcal colonized infant mice with PVM we were able to develop a novel pneumococcal-PVM model and elucidate the host, viral and bacterial dependent effects of coinfection in the upper respiratory tract.

## RESULTS

### Development of an infant mouse model of pneumococcal-PVM co-infection

To determine whether PVM infection could be established in the upper respiratory tract of infant mice, 9 day old C57BL/6 pups were given 10 plaque forming units (PFU) of PVM via intranasal administration without anaesthesia. Previously, we have shown that this dose induces a largely asymptomatic infection of the lungs when delivered by intranasal administration under anaesthesia [34,35]. Viral replication in the upper respiratory tract was measured using quantitative RT-PCR and peaked at 8 days post-infection (Figure 1A). After demonstrating PVM replication in the upper respiratory tract, we adapted our model to incorporate pneumococcal co-infection. Two pneumococcal strains were used in this study: EF3030 and BCH19. These are both clinical isolates of the same capsular serotype (19F). For our co-infection model (Figure 1B), mice were given 2×10^3^ colony forming units (CFU) of pneumococci by intranasal administration at 5 days old, followed 4 days later with PVM infection.

**Figure 1.**
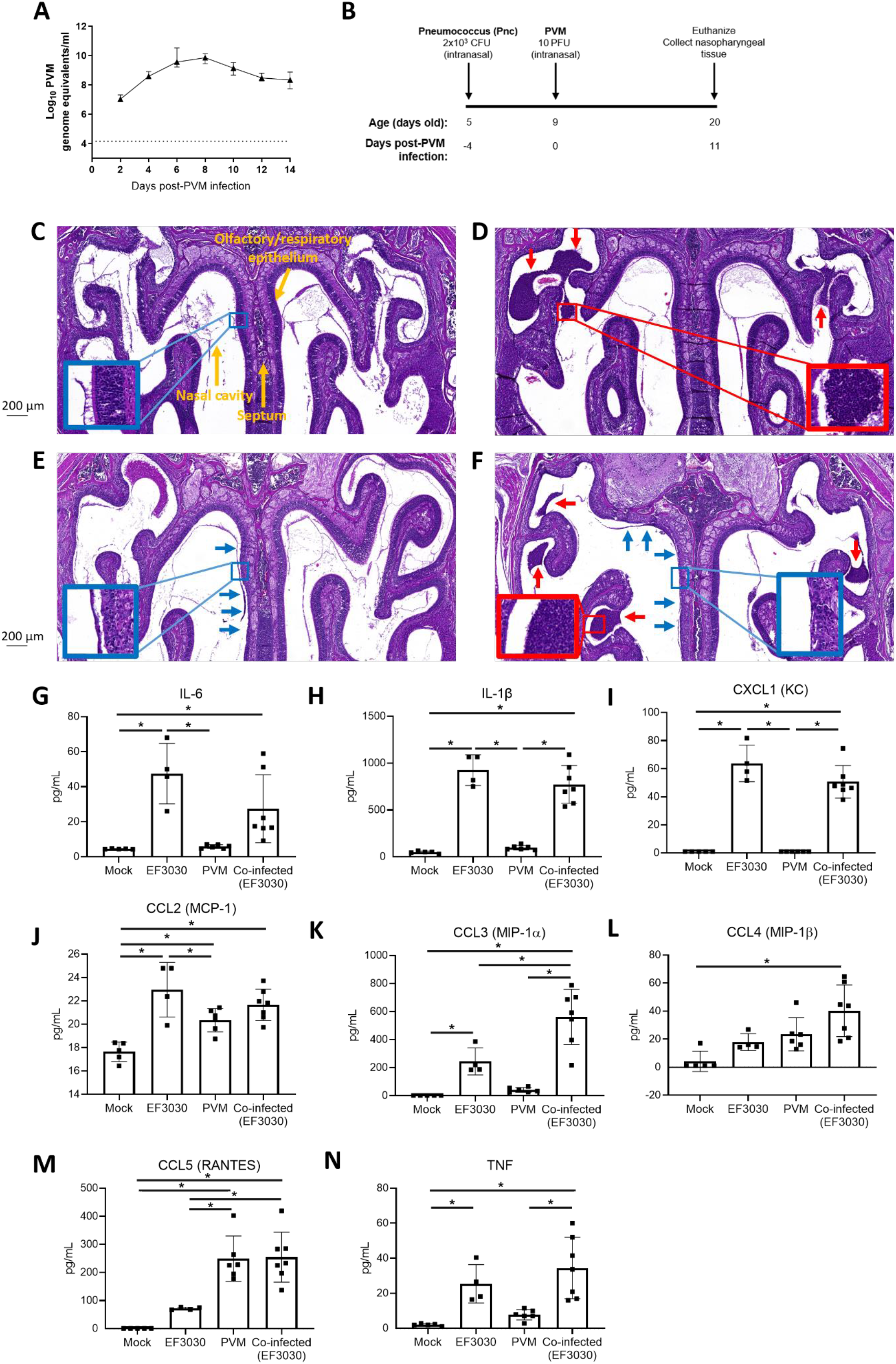
Pneumococcal-PVM co-infection model. (A) Kinetics of PVM in the nasopharynx of infant C57BL/6 mice as determined by qRT-PCR. Data are presented as the median ± IQR. (B) Schematic of pneumococcal-PVM co-infection model. PBS was administered as a vehicle control. All experiments follow this model of primary pneumococcal and secondary PVM, administration (unless otherwise stated). (C-F) Representative images of Hematoxylin and Eosin stains of nasal sections (5 μm thick). Sections were taken from 20 day old mice, who were administered (C) PBS (mock), (D) pneumococci only, (E) PVM only or (F) co-infected. Image taken using 5x magnification lens, scale bar = 200μm. Insets (denoted by boxes) are x20 magnification. Blue arrows and boxes highlight areas of necrosis along the respiratory epithelium. Red arrows and boxes highlight areas containing neutrophils. (G-N) Concentrations of pro-inflammatory cytokines in nasopharyngeal homogenates of 20 day old mice that were mock-infected, given pneumococci only, PVM only, or co-infected. Data are presented as the mean ± standard deviation and analysed by one-way ANOVA. Only p-values < 0.05 (*) are shown.

Histopathological analysis of nasal sections found that mock infected mice had no observable histopathological changes (median histopathology score of 0, Figure 1C). In contrast, mice that received pneumococci alone exhibited submucosal inflammation of the respiratory epithelium and inflammatory infiltrates (neutrophils) (median histopathology score of 3.94 [interquartile range, IQR 3.16-5.47], Figure 1D). Mice infected with PVM alone had mild submucosal inflammation, as well as unilateral olfactory sensory neuronal necrosis along the respiratory epithelium (median histopathology score of 4.5 [IQR 1.56-5.84], Figure 1E). Co-infected mice exhibited the combined effects of inflammation with neutrophils in the nasal cavity and sensory neuronal necrosis (median histopathology score of 8.19 [IQR 7.13-9.19], Figure 1F). Co-infected mice had the highest histopathological grading for tissue damage and inflammation compared with all other groups (p = 0.029 for mock, pneumococcal and PVM-infected groups, Mann-Whitney test). Using the Bliss independence test [36], we found that the effects observed in the nasal tissue of coinfected mice were synergistic rather than additive (Bliss independence score of 0.25).

We next analysed the inflammatory chemokine and cytokine responses in nasopharyngeal tissue. Assessment of pro-inflammatory cytokines present in the supernatants of homogenized samples of nasopharyngeal tissue revealed mice colonized with pneumococci (EF3030 strain) had higher concentrations of IL-6, IL-1β, CXCL1, CCL2, CCL3 and TNF compared with the mock-infected control (p < 0.001 for IL-6, IL-1β, CXCL1, CCL2 and p=0.03 for CCL3 and TNF, one-way ANOVA, Figure 1G-N). Mice infected with PVM alone exhibited elevated levels of CCL2 and CCL5 compared with the mock-infected control (p = 0.02 and < 0.001, respectively, one-way ANOVA). All chemokines and cytokines detected in the tissue of co-infected mice were elevated compared with mock-infected mice (IL-6: p = 0.04, all other chemokines and cytokines: p < 0.001, one-way ANOVA). Of note, CCL3 concentrations were increased during co-infection compared with other monoinfected mice (p=0.003 and < 0.001 for mice infected with pneumococci only and PVM only, respectively) and mock-infected experimental groups (p < 0.001) (Figure 1K). This was a synergistic rather than additive effect (Bliss independence score of 0.28).

### Effects of co-infection on pneumococci

Inflammation induced by influenza virus increases pneumococcal density [12], and nasopharyngeal samples from children with RSV have higher pneumococcal loads compared with children without RSV [37,38]. Therefore, we examined the density of pneumococci in the nasopharynx of mice, with and without PVM administration. At 20 days old, co-infection increased the median nasopharyngeal density of BCH19 ~10-fold (5.30 [IQR 5.21-5.50] and 6.29 [IQR 5.85-6.48] log_10_ CFU per nasopharynx for mice administered BCH19 alone and co-infected, respectively, p < 0.0001, Mann-Whitney test). In contrast, PVM had no effect on the density of EF3030 (6.34 [IQR 6.18-6.47] and 6.43 [IQR 6.32-6.48] log_10_ CFU per nasopharynx for mice administered EF3030 alone and co-infected mice, respectively p = 0.137, Mann-Whitney test) (Figure 2A).

**Figure 2.**
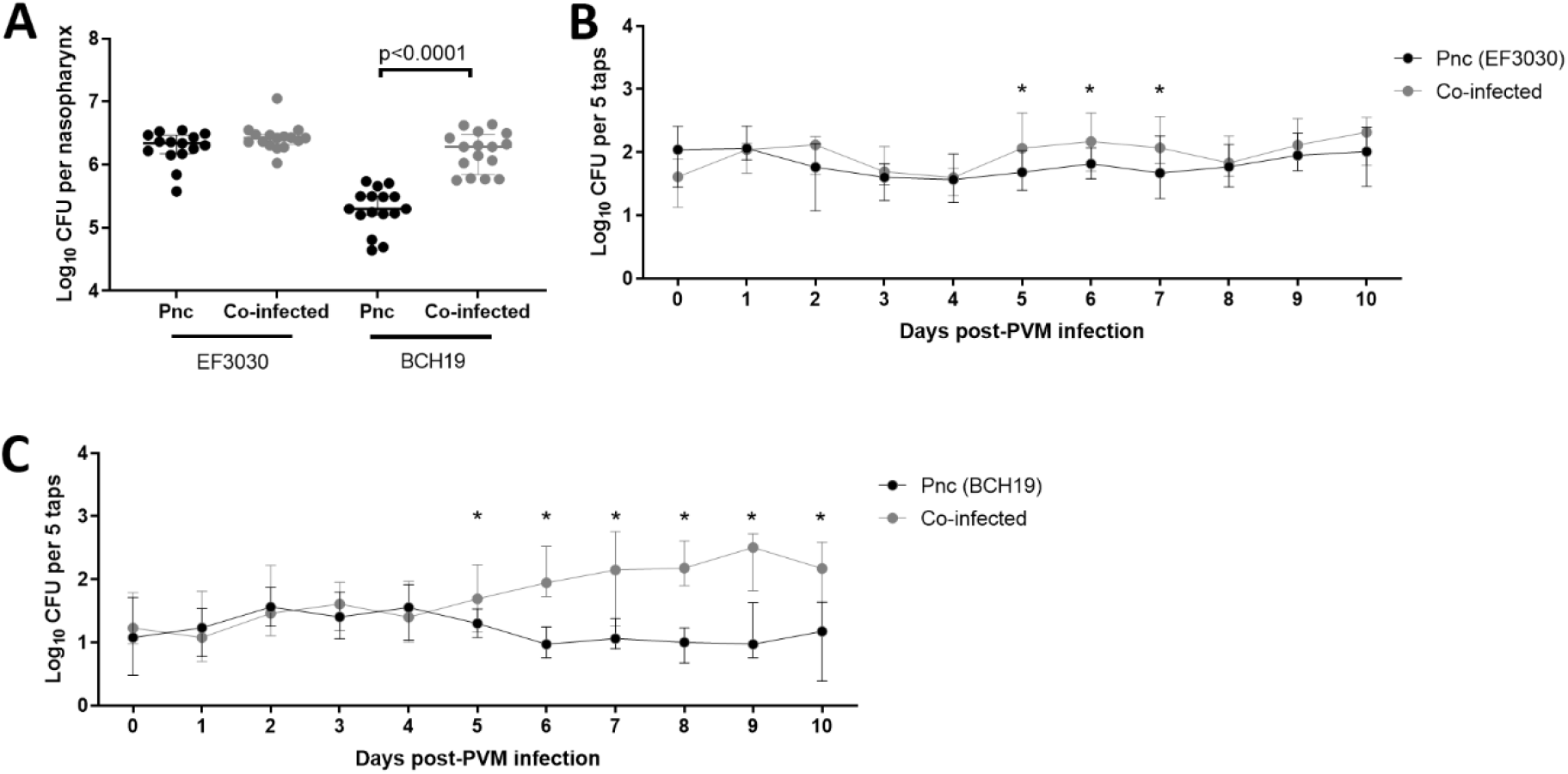
Effect of co-infection on pneumococci. (A) Nasopharyngeal density of pneumococcal strains EF3030 and BCH19 in mice given pneumococci alone (Pnc) or co-infected with PVM. (B and C) Shedding of pneumococcal strains EF3030 (B) and BCH19 (C) in the nasal secretions of co-infected mice, compared with those given pneumococci alone (Pnc). Data are presented as the median ± IQR and analysed by Mann-Whitney test. Only p-values < 0.05 (*) are shown.

Given that influenza-induced inflammation increases pneumococcal transmission [12,14], we postulated that PVM infection may have similar effects on pneumococcal colonized mice. Pneumococcal transmission involves shedding of bacteria from a colonized individual (egress), survival on fomites, and subsequently entering and colonizing a new host (acquisition) [10]. To measure egress from a colonized host, we quantified pneumococci within nasal secretions by tapping the nares of each mouse on selective media to culture pneumococci. Compared with mice colonized with pneumococci alone, mice co-infected with PVM had higher numbers of pneumococci shed in their nasal secretions from 5 days post-PVM infection (Figures 2B and 2C). Interestingly, we found that the duration and magnitude of the increase in pneumococcal shedding was dependent on the bacterial strain. PVM increased EF3030 shedding by approximately 0.36 log_10_ from 5-7 days post-PVM infection, (p = 0.006, 0.014 and 0.022, respectively Mann-Whitney test, Figure 2B). For BCH19 colonized mice, PVM increased shedding by approximately 1.04 log_10_ from 5-10 days post-PVM infection (p = 0.036, <0.0001, 0.002, <0.0001, <0.0001 and 0.0002, respectively Mann-Whitney test, Figure 2C).

To investigate acquisition, we adapted our previous pneumococcal transmission model [12], replacing influenza virus with PVM. We used the same timeline as described in Figure 1B, however, only half of each litter of mice were administered pneumococci (index mice), while the other half remained uninfected (contact mice). We then administered PVM to both index and contact mice and assessed pneumococcal colonization in contact mice 11 days post-PVM infection. This experimental design meant that PVM replication would peak in the middle of the ‘transmission window’, providing sufficient time for PVM to facilitate pneumococcal transmission. For BCH19, there was no difference in the proportion of contact mice acquiring pneumococci, when mice were co-infected with PVM (4/14, 29%) or given pneumococci alone (4/15, 27%), p = >0.999 (Fisher's exact test). For EF3030, 6/15 (40%) of contact mice acquired pneumococci when they were co-infected, compared with 2/14 (14%) of mice given pneumococci alone. However, this did not reach statistical significance (p = 0.215, Fisher’s exact test).

### Effects of co-infection on PVM

We next examined how pneumococcal colonization affects PVM. Using the same timeline for co-infection (Figure 1B), we measured shedding of PVM by swabbing the anterior nares of mice for viral quantification by qRT-PCR, every two days post-PVM administration. Mice carrying pneumococci (EF3030 strain) exhibited a 0.91 log_10_ increase in median viral shedding at 8 days post-PVM infection (6.22 [IQR 5.62-6.75] log_10_ PVM genome equivalents/ml) compared with mice given PVM alone (5.31 [IQR 4.27-6.15], p = 0.010, Mann-Whitney test, Figure 3A). This corresponds to the period of peak viral load (Figure 1A). Additionally, a trend toward increased shedding at 6 days post-PVM infection occurred, with a 0.59 log_10_ increase in shedding detected in co-infected mice (5.99 [IQR 5.14-6.72] log_10_ PVM genome equivalents/ml) compared with mice administered PVM alone (5.40 [IQR 4.97-5.95] log_1010_ PVM genome equivalents/ml), (p = 0.085, Mann-Whitney test).

**Figure 3.**
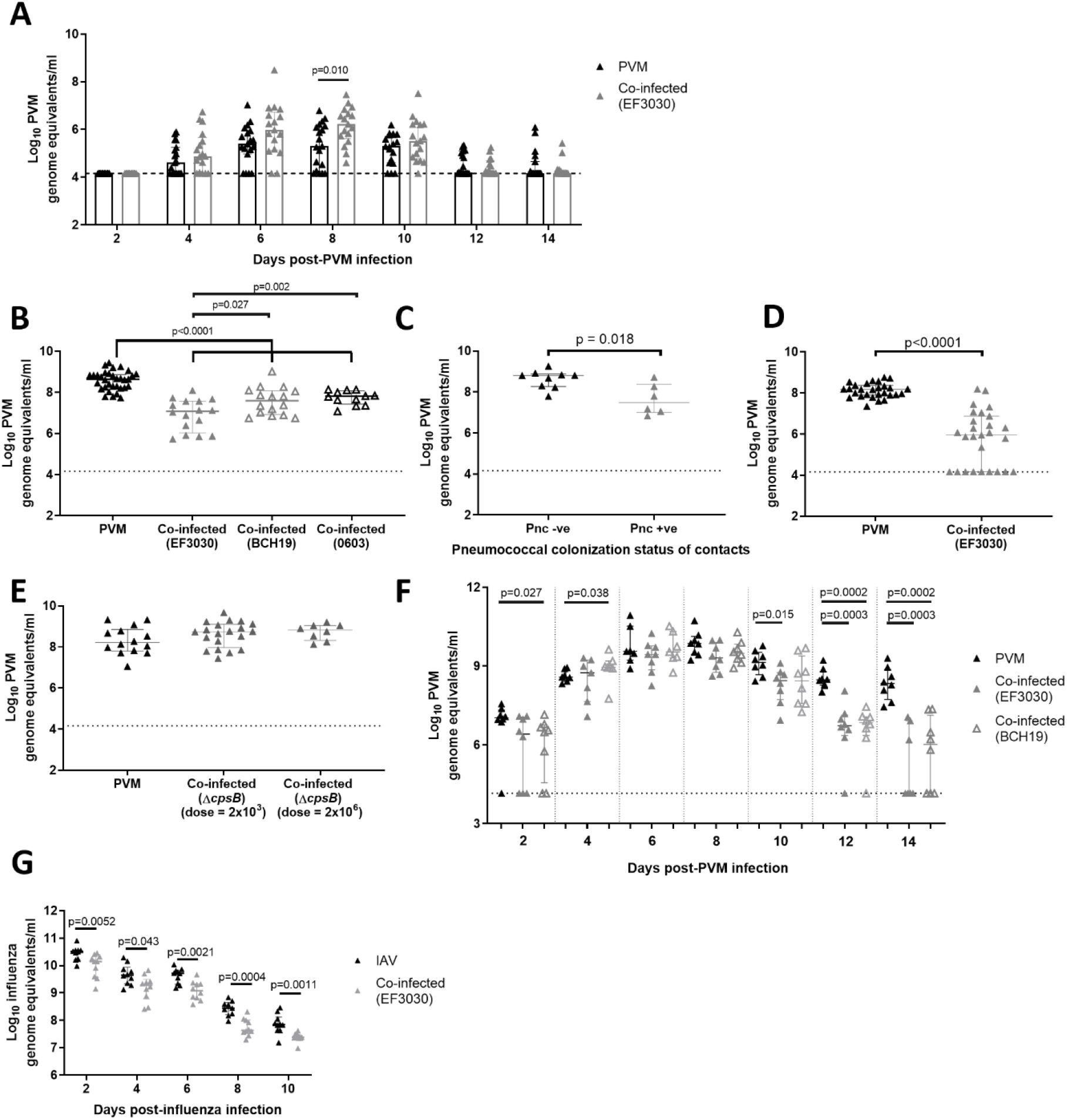
Effect of co-infection on viral infection. (A) Shedding of PVM in nasal secretions of mice co-infected with pneumococcal strain EF3030, or given PVM alone. Viral loads were determined by qRT-PCR. (B) PVM loads in 20 day old mice coinfected with pneumococci (strains EF3030, BCH19 or 0603) or given PVM alone. (C) PVM loads in the nasopharynx of 20 day old contact mice that naturally acquired pneumococci (Pnc +ve), or did not acquire pneumococci (Pnc -ve), from their co-infected littermates. Half of the litter of 5 day old pups were given pneumococcal strain EF3030 (index mice), while the other half (contacts) were not. At 9 days old, index and contact mice were given PVM. Nasopharyngeal tissue was harvested from 20 day old euthanized pups for quantification of PVM by qRT-PCR, as well as for pneumococcal culture. (D) Effect of secondary pneumococcal administration on PVM loads in 20 day old mice. Mice were given PVM at 5 days old, followed by pneumococcal administration at 9 days old. (E) PVM loads in the nasopharynx of 20 day old mice that had been co-infected with pneumococcal colonization/capsule mutant Δ*cpsB* at two doses (2×10^3^ and 2×10^6^ CFU) using the original timeline (pneumococci then PVM at 5 and 9 days old, respectively). (F) PVM loads over time in mice co-infected with pneumococci (EF3030 or BCH19), compared with mice given PVM alone. (F) influenza loads over time in mice co-infected with pneumococci (EF3030), compared with mice given influenza alone. Data are presented as the median ± IQR and analysed by Mann-Whitney test. Only p-values < 0.05 are shown.

Unexpectedly, PVM loads in the nasopharynx of co-infected mice were lower compared with those that had received PVM alone (Figure 3B). Mice colonized with EF3030 had a 1.63 log_10_ reduction in PVM loads (7.01 [IQR 6.03-7.57] log_10_ PVM genome equivalents/ml) compared with mice given PVM alone (8.64 [IQR 8.25-8.87], p <0.0001, Mann-Whitney test). Mice colonized with BCH19 had a 1.05 log_10_ reduction in PVM loads (7.59 [IQR 6.92-8.08] log_10_ PVM genome equivalents/ml) compared with mice given PVM alone (p <0.0001, Mann-Whitney test). As both EF3030 and BCH19 are serotype 19F, we tested a third pneumococcal strain (0603, serotype 6B) finding it also caused a reduction in viral loads, as there was a 0.84 log_10_ reduction in PVM loads in co-infected mice (7.81 [IQR 7.43-8.08] log_10_ PVM genome equivalents/ml) compared with mice given PVM alone, p <0.0001, Mann-Whitney test (Figure 3B).

In the transmission experiment described above, 40% of PVM-infected contact mice acquired EF3030 from their index littermates. Interestingly, when we examined PVM loads in contact mice, those that had acquired EF3030 (from their littermates) had reduced PVM loads (7.45 [IQR 7.00-8.38] log_10_ PVM genome equivalents/ml) compared with contacts that were not colonized with EF3030 (8.81 [IQR 8.27-8.88], p = 0.018, Mann-Whitney test, Figure 3C). Therefore, we hypothesized that pneumococcal-mediated antagonism of PVM may also occur when pneumococci are acquired after viral infection. To test this, we reversed the order of infection in our model by administering PVM at 5 days of age, followed by pneumococci at 9 days old. We found that the secondary pneumococcal administration also induced a reduction (2.2 log_10_) in PVM loads (5.95 [IQR 4.16-6.88] log_10_ PVM genome equivalents/ml) compared with mice given PVM alone (8.17 [IQR 7.90-8.37], p <0.0001, Mann-Whitney test, Figure 3D).

To examine the contribution of bacterial factors to the reduction of PVM loads, we generated a pneumococcal strain deficient in the ability to produce capsular polysaccharide, by deleting *cpsB* (also known as *wzh*), one of the major regulatory genes of polysaccharide synthesis. Δ*cpsB* strains produce less capsule [39,40], and this gene is important for nasopharyngeal carriage in mice [41]. Consistent with this, we were unable to recover any Δ*cpsB* mutants from the nasopharynx of 20 day old mice (n=16 mice). In contrast, the EF3030 parent strain colonized the nasopharynx of all mice at the same time point (Figure 2A). In our co-infection model, the Δ*cpsB* mutant did not reduce PVM loads (8.75 [IQR 7.97-9.10] log_10_ PVM genome equivalents/ml) compared with mice given PVM alone (8.21 [IQR 7.80-8.86], p = 0.259, Mann-Whitney test, Figure 3E). Increasing the dose of Δ*cpsB* by 1000-fold did not restore the reduction in PVM loads (8.83 [IQR 8.32-9.04], p = 0.165, Mann-Whitney test, Figure 3E).

To investigate the reduction in PVM loads in mice colonized with pneumococci further, we evaluated viral loads over time. PVM was administered to 9 day old mice, four days after they had received pneumococci. Viral loads were then measured every two days post-PVM infection. There was a significant difference in PVM loads in mice colonized with BCH19 compared with the control (PVM only) at 2 and 4 days post-PVM infection (p = 0.027 and 0.038, Mann-Whitney test), respectively (Figure 3F). Otherwise, viral loads were similar between the control and co-infected mice for the first 8 days post-PVM infection (Figure 3F). During the resolution phase (from 10-14 days post-PVM infection), all groups had a reduction in viral loads over time. Mice colonized with pneumococci had a greater reduction in PVM loads at each time point in the resolution phase, compared with controls given PVM only (Figure 3F). Additionally, the magnitude of the reduction in PVM loads in pneumococcal-colonized mice (compared with mice given PVM only) increased with each time point. There was a 0.7, 1.75 and 4.19 log_10_ reduction in PVM loads in EF3030-colonized mice at 10, 12 and 14 days post-PVM infection (p = 0.015, 0.0003 and 0.0003, Mann-Whitney test), respectively. Similarly, for BCH19-colonized mice there was a 1.62 and 2.29 log_10_ reduction at 12 and 14 days post-PVM infection (p = 0.0002 for both time points, Mann-Whitney test), respectively.

Previously, we had identified that infant mice colonized with pneumococci had reduced influenza loads in the nasopharynx [12]. We now hypothesized that pneumococcal colonization accelerates the clearance of influenza from the nasopharynx in a similar manner to its effect on PVM. To ascertain the effect of pneumococci on influenza, we administered influenza virus to 14 day old mice, 9 days after they had received pneumococci and measured influenza loads in the nasopharynx over time. Compared with mice given influenza only, mice colonized with pneumococci had reduced influenza loads at all tested time points (Figure 3G). There was a 0.37, 0.37, 0.64, 0.81 and 0.46 log_10_ reduction in influenza loads in EF3030-colonized mice at 2, 4, 6, 8 and 10 days post-influenza infection (p < 0.05 for all timepoints, Mann-Whitney test), respectively. Unlike PVM, influenza loads in co-infected mice were never comparable to mice given influenza alone.

## DISCUSSION

In this study, we have developed a novel model of pneumococcal-RSV co-infection using PVM, the murine equivalent of RSV. Our results demonstrate the complexity of bacterial-viral interactions, identifying both synergistic and antagonistic effects during co-infection. Our mouse model employs a natural cognate virus/host approach, thereby requiring lower doses and allowing robust viral replication more reflective of a natural upper respiratory tract infection [30–32]. Additionally, we use an order of infection that we hypothesize is more relevant to high-risk settings [8].

Mice that were colonized with pneumococci had inflammatory infiltrates, namely an influx of neutrophils in the nasal tissue (Figure 1C). This is consistent with previous studies in mice [42,43]. Pneumococcal infection of air-liquid interface cultures increases the production of inflammatory cytokines IL-1β and TNF [20]. Studies involving re-stimulation of human nasal cells found that those derived from participants colonized with pneumococci had elevated levels of CCL3, IL-6 and TNF [44]. Increased production of these chemokines and cytokines in response to pneumococcal colonization was also observed in our infant mouse model (Figure 1G-N). We also observed that PVM infection was associated with olfactory sensory neuronal necrosis (Figure 1E), consistent with a mouse model of RSV which showed a tropism of RSV for nasal olfactory sensory neurons and induction of necrosis along the airway epithelium [45,46], as well as evidence of neurological sequelae associated with RSV infection in humans [47–49]. Additionally, we found increased production of CCL2 and CCL5 in the nasopharynx in response to PVM infection (Figures 1J and 1M). To our knowledge, there are no studies investigating the immune response to PVM in the upper respiratory tract. However, chemokine profiles from human nasal wash samples [50,51] and mouse lungs infected with RSV [52,53] also show higher levels of CCL2 and CCL5. In our study, PVM infection increased the nasopharyngeal density of BCH19 (Figure 2A), similar to that observed in nasopharyngeal samples from RSV patients [37,38]. We also identified an enhanced inflammatory response during co-infection, supported by both histopathological analysis and pro-inflammatory cytokine and chemokine panels (Figure 1). This finding is consistent with analysis of inflammation in ciliated epithelial *ex vivo* cultures, which is augmented during pneumococcal-RSV co-infection [20].

The effects of PVM infection on the nasopharyngeal bacterial load of mice colonized with pneumococcus was dependent on the pneumococcal strain. Nasopharyngeal density of BCH19 increased during PVM infection whereas the density of EF3030 was unchanged compared to mice given pneumococci alone (Figure 2A). In the absence of PVM, infant mice carry these strains at different densities (Figure 2A). In our experience using various pneumococcal strains in infant mouse colonization models, we have not observed nasopharyngeal colonization above ~10^6^ CFU, which is consistent with other infant mouse colonization studies [13,54–56]. As such, it is plausible that ~10^6^ is the maximum density that pneumococci can reach in the nasopharynx of infant mice and thus is not altered by any changes to the microenvironment that PVM induced.

We found that PVM infection increased pneumococcal egress (Figure 2B-C), a critical phase of transmission [10], which begins as the virus is approaching peak loads (Figure 1A). The increase in pneumococcal shedding occurred with both strains but was more pronounced for BCH19, in both duration and magnitude. In the absence of viral infection, BCH19 shedding was ~1 log_10_ lower than EF3030 (Figure 2B-C). It is plausible that PVM infection has a more pronounced effect on pneumococcal strains that are usually shed in lower numbers in the absence of virus.

Influenza enhances pneumococcal transmission from index (colonized hosts) to contacts in animal models [12,14] and household contact studies [15]. In contrast, a cotton rat model of pneumococcal-RSV co-infection did not find evidence that RSV increased pneumococcal transmission [57]. However, the contacts in the cotton rat model were not infected with RSV, which at least in the influenza mouse model is important for facilitating pneumococcal acquisition [12,14]. Although we administered PVM to both index and contact mice, we found no significant increase in the acquisition of pneumococci in the presence of virus. Overall, our data suggest that, PVM infection enhances the early stages of pneumococcal transmission (egress from a colonized host) but has more limited effects on the acquisition stage.

Our study uncovered novel evidence of pneumococcal-mediated antagonism on PVM infection. Mice colonized with pneumococci had reduced PVM loads in the upper respiratory tract (Figure 3B). Examination of viral loads over time identified that this reduction was unlikely due to pneumococcal-mediated inhibition of PVM replication as viral loads in PVM-infected and co-infected mice peaked at the same time (6-8 days post-PVM infection). Some significant differences were also noted in the early stages of PVM replication where mice co-infected with BCH19 had lower and higher PVM loads compared with mice given PVM alone at 2 and 4 days post-PVM infection, respectively. However, the lack of a trend suggests these differences were due to chance. In contrast, during the resolution phase, the magnitude of the reduction increased over time. By 14 days post-PVM infection, PVM was no longer detectable in 45% of pneumococcal-colonized mice but still detectable in 100% of mice infected with PVM alone at the same time point (Figure 3F). These data demonstrate that pneumococcal colonization may create an environment capable of accelerating clearance of PVM from the nasopharynx.

This antagonistic effect of pneumococci on PVM was conserved regardless of the pneumococcal strain or serotype tested, or even the order of infection. When we infected mice with Δ*cpsB*, a mutant pneumococcal strain deficient in capsule production and incapable of carriage, this antagonistic effect was abrogated (Figure 3E), suggesting a robust immune response to pneumococcal carriage and/or capsule is required for the accelerated clearance of virus from the upper respiratory tract. In our murine co-infection model, CCL3 levels were elevated following pneumococcal colonization and this increased even further during PVM co-infection (Figure 1K). CCL3 is a chemoattractant for neutrophils, monocytes, dendritic cells and T cells [58,59]. This chemokine plays a role in the clearance of microbes such as influenza in the lung [60], murine cytomegalovirus in the liver and spleen [61], *Klebsiella pneumoniae* in the lung [62], and non-typeable *Haemophilus influenzae* in the middle ear [63]. CCL3 knockout mice have increased PVM loads in the lungs and reduced recruitment of immune cells, demonstrating this chemokine plays an important role in the control of PVM infection [64]. Future experiments will test the role of CCL3 and other effectors in pneumococcal-mediated accelerated clearance of PVM.

Adding to the complexity of these bacterial-viral interactions, we found that the timing and magnitude of accelerated PVM clearance mediated by pneumococcal colonization varied by strain. EF3030-colonized mice had a greater reduction in PVM loads at 20 days old (11 days post-PVM infection) than mice carrying BCH19 (Figure 3B), and this finding was also confirmed in our time course experiments (Figure 3F). In humans, the immune response to pneumococcal colonization positively correlates with pneumococcal density, with higher loads in the nasopharynx associated with increased levels of MMP-9, IL-6 [65] and CXCL10, as well as monocyte recruitment [44]. It is therefore plausible that the timing and magnitude of pneumococcal-mediated clearance of PVM is influenced by bacterial density, with high-density colonizing strains eliciting a greater immune response, leading to a stronger and earlier anti-PVM response.

The magnitude of the synergistic and antagonistic effects observed were dependent on the pneumococcal strain. PVM infection has a greater positive effect on BCH19 compared with EF3030 (Figures 2B and 2C). By contrast, EF3030 had a stronger and earlier negative effect on PVM compared with BCH19 (Figures 3B and 3F). As both strains are serotype 19F, these differences cannot be attributed to the pneumococcal capsule. One striking difference of these strains is the density to which they colonize the murine nasopharynx in the absence of co-infection (Figure 2A), which can influence pneumococcal shedding [54] and the immune response that may be relevant to antagonism. Future studies should focus on investigating strain-specific factors, including the role of pneumococcal density in determining the magnitude of synergistic and antagonistic outcomes of co-infection.

Lastly, we found that mice carrying pneumococci also had an antagonistic effect on influenza loads in the nasopharynx (Figure 3G). However, the reduction in influenza loads in co-infected mice compared with controls given influenza alone occurred across all time points tested. This suggests that pneumococcal colonization may act to restrict replication of influenza rather than accelerate clearance. Nonetheless, both our PVM and influenza studies demonstrate that pneumococcal colonization creates an environment in the upper respiratory tract that lowers viral loads.

Bacterial-viral interactions have important implications for vaccination strategies. For example, co-administration of whole-inactivated influenza and pneumococcal vaccines boosts protection in mice subsequently challenged with a lethal dose of influenza [66]. A randomized control trial by Harris and colleagues identified that modulating the gut microbiome using narrow-spectrum antibiotic vancomycin improved rotavirus vaccine immunogenicity [67]. More recently, a clinical trial testing BCG, a vaccine for tuberculosis, had off-target protective effects against viral respiratory infections [68]. Therefore, the pneumococcal-mediated viral antagonism observed in our study could have implications for bacterial and viral vaccines. For example, development of a pneumococcal vaccine with this antagonistic feature may also reduce viral disease. In contrast, if vaccinees carry pneumococci at the time of vaccination, the efficacy of live-attenuated viral vaccines may be reduced. Interestingly, a recent human challenge study found that the immune response to live-attenuated influenza vaccine was impaired in adults colonized with pneumococci [69], with some evidence for reduced viral loads (as inferred from qPCR Ct values) in colonized individuals.

Here, we have developed and tested a novel model of pneumococcal-RSV coinfection that is the first to show both synergistic and antagonistic interplay between these two pathogens. Taken together, our study demonstrates that the dynamics of pneumococcal-viral co-infection are more complex than initially anticipated. This study provides the basis to explore the mechanism underpinning these observations, and the implications of microbial interactions for bacterial and viral vaccination strategies.

## MATERIALS AND METHODS

### Bacterial and viral strains

Three pneumococcal strains were used in this study: EF3030 (serotype 19F, multilocus sequence type (ST) 43) [70], BCH19 (serotype 19F, ST81) and 0603 (serotype 6B, ST1536) [71]. The Δ*cpsB* mutant was constructed in a EF3030 background using genomic DNA from a previously constructed Δ*cpsB* mutant kindly provided by Dr Alistair Standish. A vial of EF3030 culture (OD 0.1) was incubated with 500 μl CTM (1% [w/v] casamino acids, 0.5% [w/v] tryptone, 0.5% [w/v] NaCl, 1% [w/v] yeast extract, 16 μM K_2_HPO_4_, 0.2% [w/v] glucose 150 μg/ml glutamine), and 55 ng CSP-1 for 10 min at 37°C with 5% CO_2_ prior to the addition of ~50 ng genomic DNA. The vial was then incubated at 32°C for 30 min and then 37°C with 5% CO_2_ for 4 h. Transformants were isolated on Horse Blood Agar (HBA) supplemented with 0.2 μg/ml erythromycin. Deletion of the *cpsB* gene was validated by whole genome sequencing. To prepare infectious stocks, strains were grown in THY broth (3% [w/v] Todd-Hewitt broth, 0.5% [w/v] yeast extract) statically at 37°C with 5% CO_2_ to an OD_600_ of ~0.4. Glycerol was added to a final concentration of 8% (v/v) and cultures were aliquoted and stored at −80°C until required. PVM strain J3666 was used in this study. Infectious stocks were prepared by purification of PVM virions from mouse lung homogenates as described previously [72]. Influenza strain A/Udorn/307/72 (H3N2) was used for this study and stocks prepared as described previously [73].

### Infant mouse model

All mouse experiments were conducted under approval by the Murdoch Children's Research Institute (MCRI) Animal Ethics Committee (A832, A857 and A898) in accordance with the Australian code for the care and use of animals for scientific purposes. C57BL/6 mice were imported from the Walter and Eliza Hall Institute and housed at the animal facility at MCRI with unlimited access to food and water. At 5 days old, pups were given 2×10^3^ CFU of pneumococci in 3 μl PBS intranasally without anaesthesia. At 9 days old, pups were administered 10 PFU of PVM in 3 μl PBS intranasally without anaesthesia. For studies involving influenza, pups were administered 20 PFU of influenza in 3 μl PBS intranasally without anaesthesia at 14 days old. Any control groups not receiving both microbes (e.g. mock or monoinfection groups) were administered PBS as a vehicle control. Pups in this model did not exhibit any overt clinical signs of illness and were housed with their dam for the duration of the experiment. Unless otherwise stated, mice were euthanized at 20 days old by cervical dislocation following anaesthesia. Nasopharyngeal tissue was harvested, placed in 1.5 ml RPMI 1640 medium (Sigma-Aldrich), and homogenized. For the enumeration of pneumococci, tissue homogenates were serially diluted and plated on HBA supplemented with 5 μg/ml gentamicin to select for pneumococci. To quantify viral loads, nasopharyngeal homogenates were subjected to centrifugation at 1400 x g for 10 min at 4°C. The supernatants were then used for viral RNA extraction and qRT-PCR.

### Pneumococcal and viral shedding

Shedding of pneumococci in the nasal secretions of mice was evaluated using a method adapted from previously described approaches [74,75]. Briefly, the nose of the mouse was tapped 5 times across two HBA plates supplemented with 5 μg/ml gentamicin. Using a 10 μl disposable loop dipped in PBS, the nasal secretions were spread across the plate. Plates were then incubated as described above, and colonies counted to calculate the CFU/5 taps.

To assess shedding of PVM in mouse nasal secretions, an infant FloqSwab (Copan) was moistened in PBS and used to swab the anterior nares of the mouse five times. The swab was then placed in 200 μl of PBS. Samples were vortexed for 5 sec and stored at −80°C until required for viral RNA extraction and qRT-PCR.

### Extraction of viral RNA, and qRT-PCR detection of PVM or influenza

Extraction of viral RNA from supernatants of nasopharyngeal homogenates or swab aliquots was performed using QIAamp Viral RNA Mini Kit (Qiagen) as per the manufacturer’s instructions, except that RNA was eluted in 30 μl of AVE buffer. Detection and quantification of viruses was conducted by qRT-PCR using the SensiFAST probe no-ROX one step kit (Bioline). The assay for detection of PVM uses primers and probe targeting the SH gene (Forward primer: 5’-ATGACCAGCAGCCGCATTGG-3’, reverse primer: 5’-TGCTTCTACTGCTGCAGGCC-3’, probe: 6FAM-CCTAACAGCTCTTCTCCTTGCATGTGC-BHQ1). The assay for detection of influenza targets the M gene using primer and probe sequences reported previously [76] (Forward primer: 5’-GGACTGCAGCGTAGACGCTT-3’, reverse primer: 5’-CATCCTGTTGTATATGAGGCCCAT-3’, probe: 6FAM-CTCAGTTATTCTGCTGGTGCACTTGCCA-BHQ1). To each 20 μl reaction containing mastermix, 0.4 μM of each primer and 0.1 μM of probe, 4 μl of viral RNA was added and run under the following cycling conditions: 45°C for 10 min, 95°C for 2 min and 40 cycles of 95°C for 15 sec and 60°C for 30 sec. Viral genome copy number was calculated using a standard curve prepared with vector DNA consisting of the SH or M gene cloned into pGEM®-T Easy (Promega) and expressed as log_10_ genome equivalents/ml of eluted RNA.

### Cytometric bead array

Nasopharyngeal homogenates were prepared as described above, except that tissues were homogenized in 500 μl RPMI, cell debris pelleted by centrifugation (1400 x g for 10 min at 4°C) and supernatants assayed for chemokine and cytokine levels using the CBA flex set (BD Bioscience) in accordance with the manufacturer’s instructions. Data were acquired using a FACSCanto II flow cytometer and concentrations calculated by comparing to standard curves generated for each chemokine and cytokine.

### Histopathological analysis of nasopharyngeal tissue

Following euthanasia, mouse heads were fixed in 10% (v/v) neutral buffered formalin and sent to the Australian Phenomics Network Histopathology and Organ Pathology Service at The University of Melbourne for sectioning and histopathological analysis. Sections (5 μm thickness) were prepared from the heads at five rostral levels at 200 μm apart. For the assessment of inflammation and tissue damage, Hematoxylin & Eosin stains were conducted on tissue sections. Samples were scored for histopathological changes. Samples were scored on a scale of 1-3 in three categories: exudate within the lumen of the nasal cavity, inflammation/number of inflammatory cells, and epithelial changes (e.g. loss of cilia, loss of epithelial cell layers). Scoring was conducted by the Australian Phenomics Network Histopathology and Organ Pathology Service, The University of Melbourne and an independent veterinary pathologist (Dr Fenella Muntz). Both operators were blinded to infection group.

### Statistical analysis

All graphs were produced and statistical analyses were conducted using GraphPad Prism version 8.4.2 (GraphPad Software). Data that were normally distributed were expressed as the mean +/- standard deviation and analysed by unpaired t test, or one-way ANOVA for with Tukey's tests for pairwise comparisons. Otherwise, data were log_10_ transformed and expressed as the median +/- IQR, and groups compared by Mann-Whitney test. Categorical data were analysed by Fisher's exact test. For all statistical analyses, an α=0.05 cut-off was used to define a significant difference between groups.

To determine if the increased effects on nasal tissue damage and CCL3 levels during co-infection were synergistic, we adapted the Bliss independence model [36], which predicts the expected combined effect of multiple stimuli. The Bliss independence model is defined as *f_xy, P_ = f_x_ + f_y_-(f_x_)(f_y_)*, where *f_x_* and *f_y_* are the effect of stimuli x and y respectively. The relationship was evaluated by comparing the experimentally observed effect of co-infection (*f_xy, O_*) to the predicted (additive) effect (*f_xy, P_*) and the Bliss independence score calculated using the formula: *f_xy, O_ - f_xy, P_*. A Bliss independence score of 0, >0 or <0 indicates an independent (additive), synergistic or antagonistic effect of co-infection, respectively.

## ACKNOWLEDGEMENTS

We thank the Translational Microbiology group at MCRI for laboratory support. We thank Professor Richard Malley for providing the BCH19 and 0603 strains used in this study. We thank Dr Alistair Standish for providing DNA used to construct the Δ*cpsB* mutant, as well as the Australian Phenomics Network Histopathology and Organ Pathology Service and Dr Fenella Muntz for histopathology and scoring services. We also thank the MCRI Disease Model Unit for services and support provided during animal studies.

## AUTHOR CONTRIBUTIONS

Conceptualization: SM, CS

Formal Analysis: SM, JM, JJ, CN

Funding Acquisition: SM, JM, KM, SP, OW, CS

Investigation: SM, JM, JJ, VW

Methodology: SM, OW, SP, CS

Resources: AU, VW, SP, OW, CS

Writing – Original Draft Preparation: SM, CS

Writing – Review & Editing: SM, JM, JJ, AU, VW, KM, OW, SP, CS

